# Evolutionary history of alpha satellite DNA in Cercopithecini: comparative cytogenomics highlights the diversification pattern of primate centromere repeats

**DOI:** 10.64898/2026.04.19.719437

**Authors:** Lauriane Cacheux, Bernard Dutrillaux, Michèle Gerbault-Seureau, Violaine Nicolas, Loïc Ponger, Bertrand Bed’Hom, Christophe Escudé

## Abstract

**Background:** Alpha satellites, a superfamily of AT-rich tandem repeats, are the primary DNA component of centromeres in Platyrrhini and Catarrhini. Analyses of the human genome suggest that centromeres behave like biological ridges, with new alpha satellite families expanding at the centromere core, splitting and displacing older ones towards the pericentromeres. The Cercopithecini tribe, which displays an unusual chromosomal evolution involving multiple chromosomal fissions and centromere formations, represents a promising model to enhance our understanding of alpha satellite DNA evolutionary history. We previously applied targeted sequencing to centromere DNA from two distant species drawn from the Cercopithecini terrestrial and arboreal lineages, and characterized six alpha satellite families exhibiting varying mean sequence identities.

**Methods:** Combining classical and molecular cytogenetics, we mapped the chromosomal distribution of these alpha satellite families across 13 Cercopithecini, one Papionini, and one Colobinae species. A nuclear marker-based phylogeny provided an evolutionary framework for interpretation.

**Results:** Our phylogeny identifies the terrestrial and arboreal lineages, and a newly designated ‘swamp clade’. We observed significant interspecies variations in alpha satellite patterns, including differences in presence/absence and distinct chromosomal distribution patterns (centromeric, pericentromeric, or subtelomeric). Families previously described as heterogeneous (83–87% mean sequence identity) exhibit a centromeric position in the swamp lineage, which is characterized by conserved karyotypes. In contrast, these families show a pericentromeric distribution in the terrestrial and arboreal lineages, replaced at the centromere core by more homogeneous families (95–98% mean sequence identity). In the arboreal clade, which is characterized by highly fissioned karyotypes, putative evolutionary new centromeres show a unique co-occurrence of highly homogeneous and heterogeneous families.

**Conclusion & Implications:** We propose a comprehensive evolutionary scenario for alpha satellite DNA in Cercopithecini, where younger families arise at the centromere core, shift toward the pericentromeres as they age, and eventually face extinction. Our study suggests that alpha satellite DNA and chromosomes evolve in an interdependent manner, with satellite diversification and displacement occurring in parallel with chromosome fissions and centromere repositioning. This comparative cytogenomic approach provides both support for the human-based evolutionary model for alpha satellite DNA and novel temporal insights into its diversification dynamics. Beyond evolutionary genomics, our findings highlight the potential of alpha satellite DNA to complement systematic studies in deciphering complex primate evolutionary histories.

## Introduction

The centromere is the specialized nucleoprotein region on eukaryotic chromosomes that coordinates the assembly of the kinetochore, thereby ensuring the faithful segregation and accurate partitioning of genetic information during mitotic and meiotic cell divisions (McKinley and Cheesman, 2016; Mellone and Fachinetti, 2021).

While the fundamental centromeric function is universally maintained through the epigenetic inheritance of the histone H3 variant CENP-A, centromeres exhibit remarkable structural and sequence variability across eukaryotes (Sullivan et al., 2001; Steiner and Henikoff, 2015). Most species possess monocentric chromosomes, but some taxa, such as certain nematodes, plants or insects, harbor holocentromeres where the region extends along the entire chromosome. Furthermore, even within monocentric organisms, centromeres can range from short, defined point centromeres supported by unique DNA sequences (e.g., *Saccharomyces cerevisiae*) to vast, complex regional centromeres composed largely of highly repetitive satellite DNA (Schueler and Sullivan, 2006; McKinley and Cheesman, 2016; Talbert and Henikoff, 2020).

The centromere region in Simians – Platyrrhini and Catarrhini – is typically composed of large arrays of tandemly repeated DNA known as alpha satellite DNA. Fundamental units, or monomers, of alpha satellite DNA are homologous AT-rich sequences of approximately 170 bp (Miga and Alexandrov, 2021). Alpha satellite DNA, having a primary function of structural scaffold for kinetochore assembly, is also actively transcribed into non-coding RNAs. These transcripts appear to play pivotal roles throughout the cell cycle by recruiting proteins for kinetochore assembly at the centromere core or by establishing and perpetuating heterochromatin within the flanking pericentromeres (McNulty and Sullivan, 2018).

During evolution, mutations and the subsequent amplification of specific monomers have collectively driven the emergence of distinct sequence families within the alpha satellite DNA. To date, over 50 alpha satellite families have been described, the majority of which have been identified in the human genome (Alves et al., 1994; Alexandrov et al., 2001; Shepelev et al., 2009; Prakhongcheep et al., 2013; Catacchio et al., 2015; Cacheux et al., 2016, 2018, Altemose et al., 2022). Owing to its highly repetitive nature, alpha satellite DNA has historically been challenging to assemble using traditional sequencing approaches, and centromeres thus constitute significant gaps in most reference primate genomes (Eichler et al., 2004; Rudd and Willard, 2004; She et al., 2004; Logsdon et al., 2024, 2025; Zhang et al., 2025). Nevertheless, the successful assembly of the boundaries of human Chromosome X centromere regions provided valuable clues about the organization of alpha satellite families within centromeres and offered critical insights into the evolutionary dynamics of alpha satellite DNA (Schueler et al., 2001; Schueler et al., 2005). Analysis of the distinct alpha satellite families within these regions resulted in the formulation of an original evolutionary model, known as the progressive proximal expansion model or gradient age hypothesis: it posits that novel alpha satellite families expand at the centromere core, consequently splitting and displacing older arrays distally toward the pericentromeric regions of each chromosome arm. The subsequent identification of several distal human alpha satellite families in the pericentromeres of other simian species corroborated this model, highlighting the need for further comparative studies to fully elucidate alpha satellite DNA evolutionary history across Primates (Shepelev et al., 2009).

The Cercopithecini, commonly known as African guenon monkeys, diverged an estimated 10 to 15 million years ago. This group currently constitutes the largest tribe within the Catarrhini infraorder, comprising more than 30 species (Guschanski et al., 2013; Kuderna et al., 2023; IUCN, 2024). Although extensive ancient and extant introgression has historically complicated their phylogenetic reconstruction (Guschanski et al., 2013; van der Valk et al., 2020; Ayoola et al., 2021; Jensen et al., 2023, 2024; Cacheux et al., 2025), nuclear data consistently support the existence of a terrestrial lineage encompassing the genera *Allochrocebus*, *Chlorocebus*, and *Erythrocebus*, and an arboreal lineage comprising the genus *Cercopithecus*. Relationships that had long remained elusive, such as the phylogenetic placements of *Allenopithecus* and *Miopithecus* (Tosi et al., 2002, 2003, 2004, 2005; Xing et al., 2007; Perelman et al., 2011, Lo Bianco et al., 2017), have been significantly clarified by recent genome-wide studies (Jensen et al., 2023, 2024). Nonetheless, some specific internal nodes within the terrestrial and arboreal clades still remain difficult to resolve definitively.

Cercopithecini exhibit a pronounced diversity of color patterns, social behaviors and ecological traits, but notably, also in chromosome morphologies (Dutrillaux et al., 1980; Glenn and Cords, 2002; Grubb et al., 2003; Enstam and Isbell, 2007; Allen et al., 2014). Indeed, comparative cytogenomics demonstrates that their karyotypes have been shaped by multiple chromosome fissions, resulting in a diploid number ranging from 48 to 72 across extant species (Dutrillaux et al., 1980, 1982; Finelli et al., 1999; Stanyon et al., 2005; Moulin et al., 2008). Species from the arboreal clade exhibit the most fissioned karyotypes, whereas those from the terrestrial clade show moderately-fissioned chromosomal patterns. Conversely, the karyotype of *Allenopithecus nigroviridis* represents the most conserved condition, as it exhibits the fewest derived chromosomal rearrangements relative to the inferred ancestral state of the tribe. These large-scale genomic rearrangements are hypothesized to be associated with the formation of Evolutionary New Centromeres (ENCs), render the Cercopithecini a compelling model for investigating the evolutionary dynamics of alpha satellite DNA (Stanyon et al., 2005; Moulin et al., 2008; Rocchi et al., 2012).

Consequently, we performed high-throughput sequencing of the alpha satellite component in two key species, *Allochrocebus solatus* (terrestrial lineage) and *Cercopithecus pogonias* (arboreal lineage) (Cacheux et al., 2016, 2018). This sequencing, combined with integrated computational and cytogenomic analyses, uncovered a previously uncharacterized diversity of alpha satellite DNA within the Cercopithecini, leading to the characterization of four distinct families (termed C1 to C4), as well as two additional (sub)families within the C1 family (termed C5 and C6). We further observed that these alpha satellite families exhibited varying mean sequence identities, with the most homogeneous families (95–98% mean sequence identity) localizing to the centromere cores and the most heterogeneous ones (83–87% mean sequence identity) at pericentromeres.

In the present study, we conducted a comparative study of alpha satellite DNA in Cercopithecini. We combined classical and molecular cytogenetic approaches to examine the presence/absence and chromosomal distribution of alpha satellite families C1 to C6 in 13 species distributed across the Cercopithecini phylogenetic tree, as well as two outgroup species. In order to interpret our results within the framework of Cercopithecini species evolution, we reconstructed a nuclear-based phylogeny to encompass our full taxonomic sampling, including species not yet covered by genome-wide sequences.

We predicted that alpha satellite DNA would follow the progressive proximal expansion model and that centromere sequence evolution would be deeply intertwined with karyotype evolution, implying that:

1. significant interspecies variations would be observed across all six alpha satellite families, including differences in presence/absence and distinct centromeric and pericentromeric chromosomal distribution patterns;
2. the observed alpha satellite distribution patterns would enable the reconstruction of a comprehensive evolutionary scenario involving the successive emergence of alpha satellite families at core centromeres, their subsequent displacement toward pericentromeres as they age, and their eventual extinction (*i.e.*, complete loss of detectable copies) in certain lineages;
3. families classified as heterogeneous or homogeneous in *A. solatus* and *C. pogonias* would reflect their early or late emergence, respectively, during Cercopithecini evolution;
4. the oldest families would be retained in a centromeric position in species with conserved karyotypes, while the newest families would be confined to species with highly fissioned karyotypes;
5. putative ENCs would show a unique pattern of alpha satellite diversity.

We also predicted that the detailed distribution of alpha satellite DNA families across the Cercopithecini phylogeny would yield novel molecular support for previously unresolved nodes, thereby helping disentangle complex primate evolutionary histories.

## Methods

### Phylogenetic Reconstruction

The phylogenetic relationships among 22 Cercopithecini species were reconstructed using sequences of nine nuclear genes already employed in primate phylogenies (ABCA1, BRCA2, CFTR, DENND5A, ERC2, LRPPRC-169, SRY, TTR, and ZFX) (Perelman et al., 2011). The species included were: *Allenopithecus nigroviridis*, *Allochrocebus lhoesti*, *A. preussi*, *A. solatus*, *Cercopithecus ascanius*, *C. albogularis*, *C. campbelli*, *C. cephus*, *C. diana*, *C. erythrotis*, *C. hamlyni*, *C. mitis*, *C. mona*, *C. nictitans*, *C. petaurista*, *C. pogonias*, *C. roloway*, *C. wolfi*, *Chlorocebus sabaeus*, *C. aethiops*, *Erythrocebus patas*, and *Miopithecus ogouensis*.

#### Sample Collection and Molecular Procedures

When not retrieved from GenBank, the sequences were obtained from fibroblast samples provided by the “Tissues and Cryopreserved Cells from Vertebrates” collection (TCCV, RBCell), held by the National Museum of Natural History (MNHN) in Paris, France. These fibroblast samples were originally amplified from skin biopsies collected on captive individuals (zoological parks, rehabilitation centers), either during veterinary health checks under anesthesia or during post-mortem veterinary autopsies. DNA extraction was performed on these samples using the QIAGEN QIAamp DNA Mini Kit. The target genes were amplified using Polymerase Chain Reaction (PCR) with primers provided in Table S1. The PCR protocol consisted of 40 cycles: 30 s at 94°C (denaturation), 40 s at 57 °C (ABCA1) or 63 °C (ZFX) or 65 °C (BRCA2, LRPPRC-169, TTR) or 67 °C (CFTR, DENND5A, ERC2) (annealing), and 1 min 20 s at 72 °C (extension). The resulting double-stranded PCR products were purified and sequenced by Eurofins Genomics (Germany). All GenBank accession numbers, including those retrieved from the public database and those corresponding to the newly generated sequences for this study, are provided in Table S2.

#### Phylogenetic Inference and Model Selection

Sequences were aligned using MUSCLE (Edgar, 2004). Prior to phylogenetic inference, the computer program PartitionFinder 2 (Lanfear et al., 2016) was used to evaluate the fit of 24 models of nucleotide substitution to the data, restricted to those implemented in MrBayes 3.1 (Huelsenbeck and Ronquist, 2001). The selection process employed the greedy algorithm (Lanfear et al., 2012) with likelihoods calculated in PhyML 3.0 (Guindon et al., 2010). Initial partitions were defined by gene, and the optimal evolutionary models were selected according to the Bayesian Information Criterion (BIC). The HKY model was chosen for DENND5A and SRY, while the HKY+I model was selected for all other loci (ABCA1, BRCA2, CFTR, ERC2, LRPPRC-169, TTR, and ZFX). These models were subsequently employed in the Bayesian analysis. The phylogenetic tree was constructed using Bayesian Markov Chain Monte Carlo (MCMC) analysis. The analysis utilized two independent runs, each initialized with random trees and conducted with three heated chains and a single cold chain. A total of 25 million generations were performed per run, with trees and parameters sampled every 1000 generations. The first 10% of the sampled trees were discarded as burn-in. Convergence was confirmed by ensuring that the average standard deviation of split frequencies (ASDSF) dropped well below 0.01 (final ASDSF = 0.0022). Bayesian posterior probabilities (PP) were used to assess branch support of the MCMC tree. The phylogenetic tree was rooted using *Colobus guereza* (Colobinae), with three Papionini species (*Cercocebus torquatus*, *Macaca sylvanus*, and *Mandrillus sphinx*) also serving as outgroups to Cercopithecini (Perelman et al., 2011; Guschanski et al., 2013). All input files and configuration parameters are available in the Zenodo repository (https://doi.org/10.5281/zenodo.19650370).

### Cell Cultures and Metaphase Preparations

Fibroblast samples from one Papionini species (*Macaca sylvanus*), one Colobinae species (*Colobus angolensis*), and 13 Cercopithecini species were used for cytogenomic analysis. The Cercopithecini species included were: *Allenopithecus nigroviridis*, *Allochrocebus lhoesti*, *A. solatus*, *Cercopithecus ascanius*, *C. cephus*, *C. diana*, *C. erythrotis*, *C. mitis*, *C. mona*, *C. nictitans*, *C. pogonias*, *C. roloway*, and *Erythrocebus patas*. These samples were obtained from the TCCV, RBCell collection (MNHN, Paris). The identification numbers of the specimens used are available in Table S3.

Cell cultures and metaphase preparations were performed according to Moulin et al. (2008). Briefly, fibroblast cultures were grown at 37 °C in Dulbecco’s modified eagle medium (Gibco), supplemented with 10% fetal bovine serum (FBS, Cytiva), 2% AmnioMAX (Gibco) and 0.2X antibiotic/antimycotic solution (Cytiva). The cultures were synchronized using fluorodeoxyuridine (0.06 μg/ml, Sigma) overnight. Bromodeoxyuridine (0.02 mg/ml, Sigma) was then added for the last 8 h of culture to incorporate into late-replicating DNA. Colchicine (0.04 μg/ml, Sigma) was added for the last 3 h of culture to arrest the cell cycle at metaphase. A hypotonic shock was performed using 15% FBS and 181 μg/ml KCl for 10 min at 37 °C. The cells were subsequently fixed in a solution of acetic acid (1 volume) and ethanol (3 volumes), then spread on slides that were dried and stored at − 20 °C.

### Fluorescence *in situ* Hybridization (FISH)

FISH experiments were carried out on metaphase chromosome preparations following the methodology previously established by Cacheux et al. (2016, 2018). The hybridization utilized short oligonucleotide probes previously described in the same studies. These probes were specifically designed to target the C1 to C6 alpha satellite families identified in the *A. solatus* and/or *C. pogonias* genomes. Probe sequences are detailed in Table S4, which also specifies the positions of Locked Nucleic Acid (LNA) modifications. The specific sets of probes used for the detection of alpha satellite families in each studied species are summarized in Table S5. The LNA-modified probes were custom-synthesized and purchased from Eurogentec (Seraing, Belgium). The hybridization solutions were prepared by diluting the oligonucleotide probes to a final concentration of 0.1 µM in a buffer composed of 2X SSC (pH 6.3), 50% deionized formamide, 1X Denhardt solution, 10% dextran sulfate, 40 mM NaH_2_PO_4_, and 0.1% SDS. A 20 µL volume of the hybridization solution was deposited onto each slide and covered with a coverslip. The slides were subsequently denatured by heating at 70 °C for 3 min, followed by hybridization at 37 °C for 1 h in the Thermobrite apparatus (Leica Biosystems).

Post-hybridization washes consisted of two washes in 2X SSC at 63 °C. An occasional higher stringency wash at 68 °C was sometimes performed (see Results). Preparations were then incubated in a blocking solution (4% bovine serum albumin (BSA), 1X PBS, 0.05% Tween 20) for 30 min at 37 °C to minimize non-specific binding. Depending on the specific probe combination, the following fluorescent conjugates were used for signal detection, all diluted 1:200 (v/v) in the blocking solution (1X PBS, 0.05% Tween 20, 4% BSA) and incubated for 30 min at 37 °C: Alexa 488-conjugated streptavidin (Life Technologies, 5 µg/ml final concentration); Cy5-conjugated streptavidin (Caltag Laboratories, 1 µg/ml final concentration); FITC-conjugated sheep anti-digoxigenin (Roche, 1 µg/ml final concentration); and rhodamine-conjugated sheep anti-digoxigenin (Roche, 1 µg/ml final concentration). All subsequent washings were performed in 2X SSC supplemented with 0.05% Tween 20. Chromosomes were counterstained in blue using DAPI (4 µg/ml) and slides were mounted with Vectashield Antifade Mounting Medium (Vector Laboratories). Alternatively, chromosomes were counterstained with propidium iodide (PI, 2 µg/ml). Slides were then mounted using a p-phenylenediamine (PPD) antifade solution (1 mg/ml, pH 11), prepared by dissolving 50 mg of PPD in a mixture of 90% glycerol and 10% 1X PBS.

### Image Acquisition and Analysis

FISH observations were performed using two different epifluorescence microscope systems: a DMRB (Leica Microsystems) and an Axio Observer Z1 (Zeiss). Images were captured using cooled CCD cameras (ProgRes MFcool, Jenoptik and ORCA R2, Hamamatsu) mounted on the respective microscopes. For each individual and each FISH experiment, a minimum of 20 metaphase spreads were visualized, all of which confirmed the hybridization patterns described in the Results section. Color-combined images were reconstructed using ImageJ (Abràmoff et al., 2004), and metaphases were karyotyped using Isis 5.3 software (Metasystems, Altussheim, Germany).

### Comparative Chromosomal Mapping

To pursue the detailed chromosomal mapping of alpha satellite families, six phylogenetically dispersed Cercopithecini species were selected: *A. nigroviridis*, *E. patas*, *A. solatus*, *C. roloway*, *C. cephus* and *C. pogonias*. These species were chosen to represent the diversity of karyotype structures and the overall alpha satellite family distribution identified within the clade. This investigation focused on precisely mapping the distribution of the C2, C5, and C6 alpha satellite families.

The alpha satellite distribution profiles were compared by performing a site-by-site analysis across homologous chromosomes. The visualization of these distributions used completed karyotypes derived from the FISH experiments. The comparative mapping relied on the alignment of homologous chromosomes from the six studied species with the presumed ancestral Cercopithecidae karyotype. This alignment was established using previously published R banding patterns and chromosome painting data (Dutrillaux et al. 1982, Moulin et al. 2008).

Interpretive criteria were applied to the visualization maps to ensure an accurate representation of the relevant differences during the comparative analysis. For the C2 family, where signals were detected in varying intensity across most centromeric regions: only centro- or pericentromeric loci displaying strong C2 signals were illustrated on the map. In contrast, all telomeric or interstitial loci exhibiting C2 signals – which represent non-standard localizations – were systematically included in the visual representation for comparison.

## Results

### Cercopithecini phylogenetic relationships

Our reconstructed Cercopithecini phylogeny (Fig. 1) resolved three distinct primary lineages: the established terrestrial and arboreal clades, and a novel third lineage including the genera *Allenopithecus* and *Miopithecus*. All three primary lineages are robustly supported (PP = 1). Notably, the node defining the monophyly of the joint terrestrial and arboreal lineages exhibits moderate support (PP = 0.70).

Based on the shared ecological affinity for hydric environments of *Allenopithecus* and *Miopithecus* species, we named the newly defined lineage the ‘swamp clade’. This designation reflects the strong dependency of both taxa on water-dominated habitats: the *Allenopithecus* species (Allen’s swamp monkey) is strictly confined to lowland swamp forests, while *Miopithecus* species are strictly riverine, inhabiting gallery forests and swampy areas adjacent to major waterways (McGraw, 1994; Gautier-Hion et al., 2005; Maisels et al., 2006; Kingdon, 2015).

Within the terrestrial lineage, nodes corresponding to the genera *Chlorocebus* (*C. aethiops*, *C. sabaeus*) and *Allochrocebus* (*A. lhoesti*, *A. preussi*, *A. solatus*) are well supported (PP = 1). Similarly, strong support (PP = 1 or > 0.98) is observed within the arboreal lineage for the *Cercopithecus diana* group (*C. diana*, *C. roloway*), the *C. mona* group (*C. mona*, *C. campbelli*, *C. pogonias*, *C. wolfi*), the *C. cephus* group (*C. cephus*, *C. ascanius*, *C. erythrotis*, *C. petaurista*) and the node corresponding to the joint *C. cephus* and *C. mitis* (*C. mitis*, *C. albogularis*, *C. nictitans*) groups.

**Figure 1.**
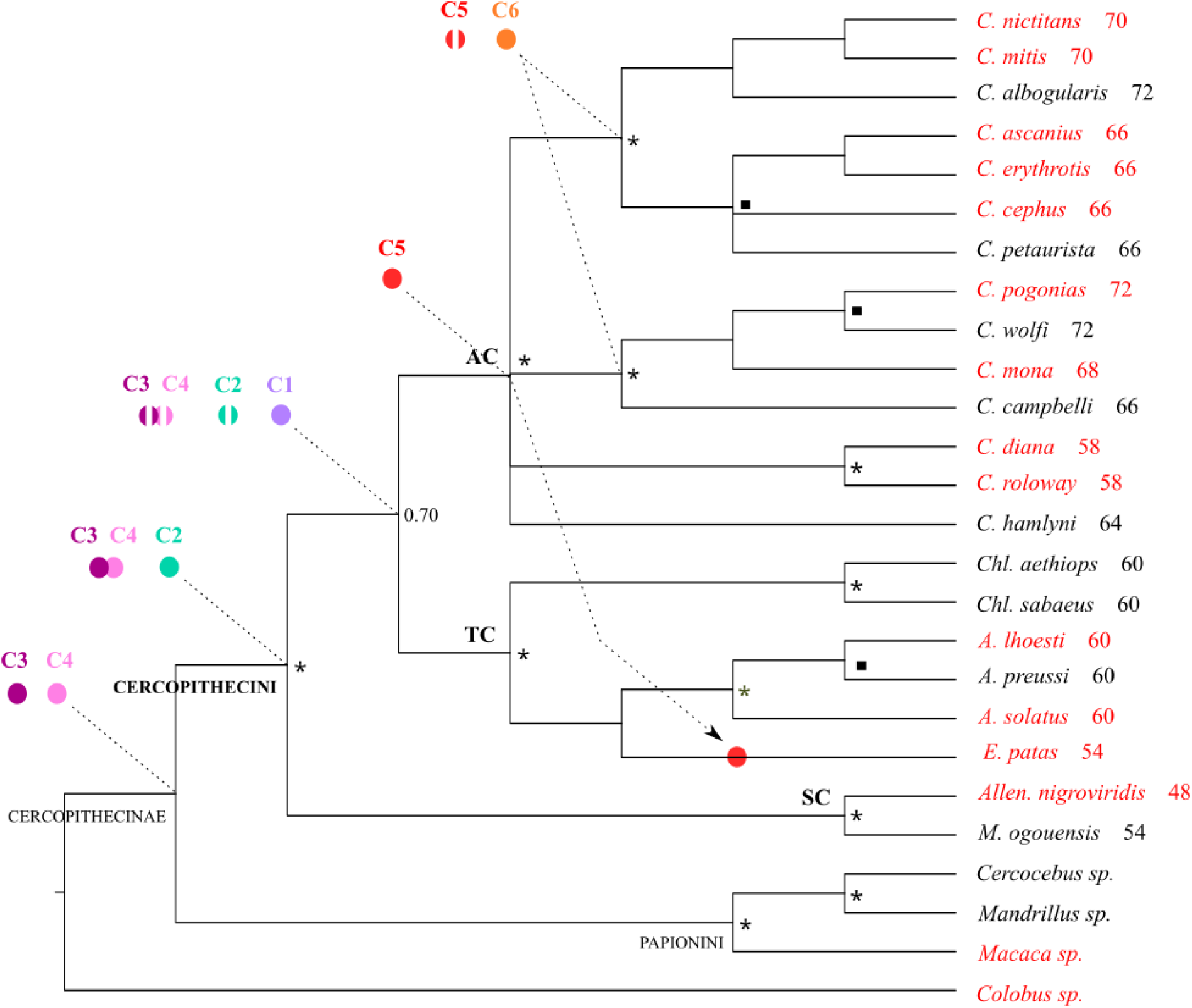
Cercopithecini phylogenetic relationships (Bayesian inference) and alpha satellite DNA evolutionary scenario. The relationships are shown as a cladogram inferred from nine nuclear markers (ABCA1, BRCA2, CFTR, DENND5A, ERC2, LRPPRC-169, SRY, TTR, and ZFX). **Taxon Designations**: The Cercopithecini primary clades and genera are labeled on the phylogeny: AC: Arboreal clade, TC: Terrestrial clade, SC: Swamp clade; *C*: *Cercopithecus*, *Chl*: *Chlorocebus*, *A*: *Allochrocebus*, *E*: *Erythrocebus*, *Allen*: *Allenopithecus*, *M*: *Miopithecus*. Numbers associated with species names represent their diploid chromosome number (2n). Species involved in the present study of alpha satellite DNA are colored in red. **PP**: Nodes with maximum PP (1.00) are labeled with a star (★) and nodes with PP > 0.98 and &< 1 are labeled with a square (▪). **Alpha Satellite Evolution**: The hypothesized emergence of the C1 to C6 alpha satellite families, inferred from FISH and sequence results, is indicated by colored circles (C1: purple, C2: pastel green, C3: dark pink, C4: light pink, C5: red, C6: orange). The displacement of certain families toward pericentromeres is indicated by broken circles. An arrow indicates hypothesized genetic transfer.

### Diversity and distribution of alpha satellite DNA across Cercopithecini genomes

Species highlighted in red on Figure 1 correspond to those whose alpha satellite DNA component was investigated by FISH. They were chosen across the three primary clades (swamp, terrestrial, and arboreal lineages) to represent the variety of karyotypes within Cercopithecini. The FISH results pertaining to the alpha satellite DNA families are summarized in Table 1.

#### Terrestrial clade of Cercopithecini

Data on the alpha satellite families present in *A. solatus* (2n=60) were previously described (Cacheux et al., 2016). Targeted sequencing of alpha satellites identified four distinct families: C1 and C2 were *in silico* characterized from a monomer dataset, while C3 and C4 were identified from a dimer dataset, the latter two systematically associating within dimeric sequences. The C1 family was found to be homogeneous, exhibiting an average sequence identity of 95%, whereas the other families, C2 (85%), C3 (86%), and C4 (83%), were comparatively more heterogeneous. FISH data subsequently revealed distinct chromosomal locations: C1 localized exclusively to core centromeres, and C2 to pericentromeres. In contrast, C3 and C4 probes showed clear signals on the Y-chromosome while yielding only faint, difficult-to-detect signals on the pericentromeres of other chromosomes (Cacheux et al., 2016). This investigation of the terrestrial clade was extended here to two additional species: *E. patas* (2n=54) and *A. lhoesti* (2n=60) (Fig. 1).

In *A. lhoesti*, C1 and C2 probe signals were observed on the centromeres (primary constrictions) and pericentromeres (regions surrounding the primary constrictions), respectively (Fig. 2A). Notably, strong C2 signals were detected on the short arms of acrocentric chromosomes. The distribution pattern of the C1 and C2 probes was broadly similar in *E. patas* (Fig. 2B, D), though this species displayed several additional, specific features: the C2 probes hybridized on both telomeric and interstitial regions of certain submetacentric chromosomes; the C1 and C2 probes co-localized on one specific chromosome pair; and the C1 probes provided notably large signals, extending toward pericentromeres, on some other chromosomes.

**Table 1.** FISH detection and chromosomal mapping of alpha satellite families (C1–C6) in Cercopithecini genomes.

|  |  | C1 | C2 | C3 | C4 | C5 | C6 |
| --- | --- | --- | --- | --- | --- | --- | --- |
| AC | *C. mitis* | + c, pc | + pc | (-) | (-) | + c, pc | + c |
|  | *C. nictitans* | + c, pc | + pc | (-) | + pc | + c, pc | + c |
|  | *C. ascanius* | + c, pc | + pc | (-) | + pc | + c, pc | + c |
|  | *C. erythrotis* | + c, pc | + pc | (-) | + pc | + c, pc | + c |
|  | *C. cephus* | + c, pc | + pc | (-) | + pc | + c, pc | + c |
|  | *C. pogonias* | + c, pc | + pc | (-) | + pc | + c, pc | + c |
|  | *C. mona* | + c, pc | + pc | (-) | (-) | + c, pc | + c |
|  | *C. diana* | + c | + pc | (-) | + pc | + c | - |
|  | *C. roloway* | + c | + pc | (-) | + pc | + c | - |
| TC | *A. lhoesti* | + c | + pc | + pc | + pc | (+) | - |
|  | *A. solatus* | + c | + pc | + pc | + pc | (+) | - |
|  | *E. patas* | + c, pc | + pc | (-) | + pc | + c | - |
| SC | *A. nigroviridis* | (+) | + c | + c | + c | (+) | - |
| Out | *M. sylvanus* | - | (+) | + na c | + na c | (+) | (+) |
|  | *C. angolensis* | - | - | - | - | (+) | - |

Notes:
**Clade Designations:** AC: Arboreal clade. TC: Terrestrial clade. SC: Swamp clade. Out: Outgroup species.
**FISH Signal Symbols:** +: Signal detected. -: Signal not detected. (-): Qualified absence. FISH signal was not detected, but other genomic data suggest the presence of residual sequences (Details in Results). (+): Non-specific signal. This signal is detected at the centromeric region but fails to meet specific criteria for structural organization or hybridization stringency (Details in Results).
**Major Chromosomal Localizations:** c: Centromeric localization (signal restricted to the primary constriction). pc: Pericentromeric localization (signal situated within the flanking regions). c, pc: Centromeric and pericentromeric localization (signal spanning the primary constriction and extending into the adjacent flanking regions).
**Structural Organization Note:** + na (not associated): Signals were detected, but the characteristic colocalization of C3 and C4 signals, which is indicative of a dimeric C3–C4 organization, was not observed.
**Source of Data:** Data pertaining to *A. solatus* (C1–C4) and *C. pogonias* (C1, C2, C5 and C6) are sourced from previously published results (Cacheux et al., 2016, 2018).

**Figure 2.**
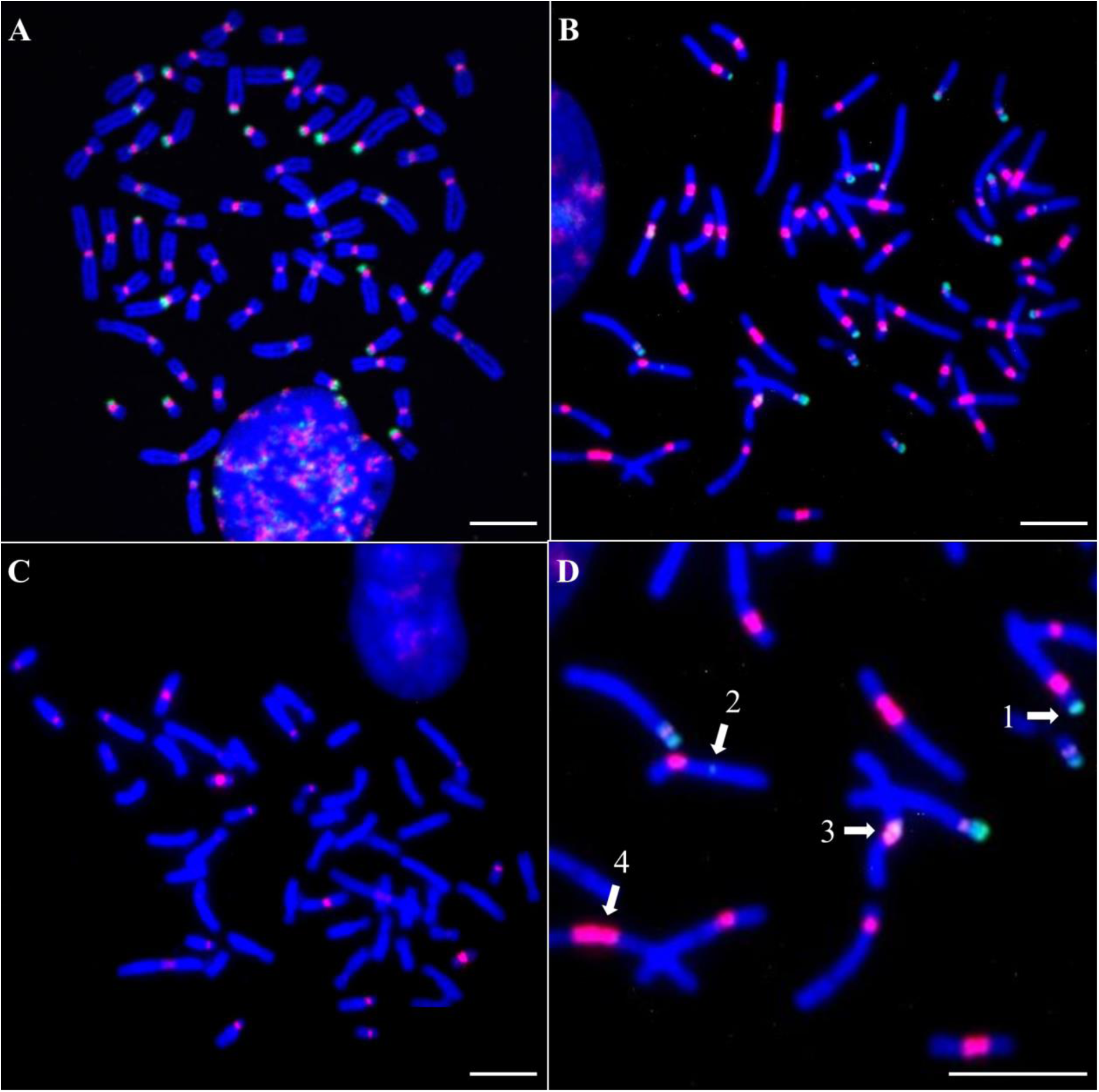
FISH analysis of alpha satellite DNA in species from the terrestrial clade. Metaphase chromosomes are shown in blue (DAPI counterstain). (A) Hybridization signal of probes C1b (red) and C2b (green) on *A. lhoesti* chromosomes. (B) Hybridization signal of probes C1a (red) and C2a (green) on *E. patas* chromosomes. (C) Hybridization signal of probe C5a (red) on *E. patas* chromosomes. (D) Magnified view of image (B), illustrating specific probe localizations in *E. patas*: 1: C2a signal labeling a telomeric region on a submetacentric chromosome, 2: C2a signal labeling an interstitial locus, 3: Co-localization of C1a and C2a signals (appearing yellow) on a centromeric region, 4: C1a signal labeling a large (peri)centromeric array. Scale bar = 10µm.

C3 probes failed to yield detectable signals on the chromosomes of *E. patas*. In contrast, C4 probes yielded faint signals on the pericentromeres of both *E. patas* and *A. lhoesti*, as did the C3 probes for the latter (Fig. S1).

While C5 probes initially produced signals on *A. lhoesti* chromosomes, the signal intensity decreased substantially when the stringent post-hybridization wash was performed at 68 °C, compared to the standard temperature of 63 °C (Fig. S2). The reliability of this stringent threshold was demonstrated by the results in *E. patas*, where C5 probes provided consistently temperature-resistant signals in the core centromeres of nine chromosome pairs (Fig. 2C and Fig. S2). This differential stability under identical high-stringency conditions indicates that the hybridization in *A. lhoesti* was likely non-specific, suggesting that the C5 family is effectively absent from its genome.

C6 probes did not yield any detectable signals on the chromosomes of either *A. lhoesti* or *E. patas*. *Arboreal clade of Cercopithecini*

Data on the alpha satellite families present in *C. pogonias* (2n=72) were previously described (Cacheux et al., 2018). Targeted sequencing analysis of monomeric alpha satellites led to the characterization of two distinct families: C1 and C2, in addition to two subfamilies within C1: C5 and C6. Consistent with the previous observation, the C1 family was found to be homogeneous (95% average sequence identity) compared to the more heterogeneous C2 (85%). The C5 and C6 subfamilies also exhibited high homogeneity, showing average sequence identities of 95% and 98%, respectively. Subsequent FISH analysis revealed a clear partitioning of these sequences: the C1 family localized at the core centromere of all chromosomes and additionally extended toward the pericentromeric regions, where it was still surrounded by the C2 family. Furthermore, the C5 subfamily displayed a variable centromeric to pericentromeric localization across several chromosomes, whereas the C6 subfamily mapped exclusively to the core centromeres, predominantly on acrocentric chromosomes.

We completed this description by analyzing a dimer alpha satellite sequencing dataset, which revealed the presence of associated C3 and C4 heterogeneous sequences in *C. pogonias*, mirroring the pattern observed in *A. solatus* (Figs. S3 and S4, Tables S6 and S7). However, in *C. pogonias*, C3 probes failed to yield detectable FISH signals, and C4 probes yielded only faint, difficult-to-detect signals on the pericentromeres (Fig. S5).

The investigation of the arboreal clade was extended here to eight additional species: *C. roloway* (2n=58), *C. diana* (2n=58), *C. mona* (2n=68), *C. cephus* (2n=66), *C. erythrotis* (2n=66), *C. ascanius* (2n=66), *C. mitis* (2n=70), and *C. nictitans* (2n=70) (Fig. 1). *Cercopithecus roloway* and *C. cephus* were chosen as representative examples to display the alpha satellite distribution in Figures 3 and 4.

For all species in the arboreal clade, C1 and C2 probe signals typically exhibited centromeric and pericentromeric localization, respectively (Fig. 3). In species displaying highly fissioned karyotypes (*C. mona*, *C. cephus*, *C. erythrotis*, *C. ascanius*, *C. mitis* and *C. nictitans*), the C1 signals additionally extended toward the pericentromeric regions (Fig. 3B, E). We observed a rare exception to the regular localization pattern in one chromosome pair of *C. roloway*, where C2 signals localized to the centromere while C1 signals extended toward the pericentromeric region (Fig. 3D). In addition to standard pericentromeric localization, C2 signals were commonly detected in some telomeric regions across all analyzed species; furthermore, C2 signals exhibited interstitial localization in *C. cephus* (Fig. 3C, E). Neither C1 nor C2 signal was detected on any Y chromosome, and only one other chromosome (an autosome in *C. cephus*) displayed a complete absence of signal for both families (Fig. 3B).

C3 probes failed to yield detectable signals on the chromosomes of any species in the arboreal clade. C4 probes yielded only faint, difficult-to-detect signals on the pericentromeres of all species, with the exception of *C. mitis* and *C. mona* where no signal was detected. This consistent pattern of extremely low or absent hybridization signals across the terrestrial and arboreal clades is put into perspective by sequencing data, which reveal that C3 and C4 families are present and associate to form dimeric C3-C4 sequences within the *A. solatus* (Cacheux et al., 2016) and *C. pogonias* genomes (Figs. S3 and S4; Table S6). The presence of these dimeric structures in such phylogenetically divergent species indicates that this genomic organization should be a shared feature across terrestrial and arboreal lineages. Consequently, the frequent absence of C4 or especially C3 signals is likely attributable to the low abundance of these dimers, falling below the FISH detection limit. Furthermore, the differential detection between the two families probably reflects inherent differences in probe hybridization efficiency.

**Figure 3.**
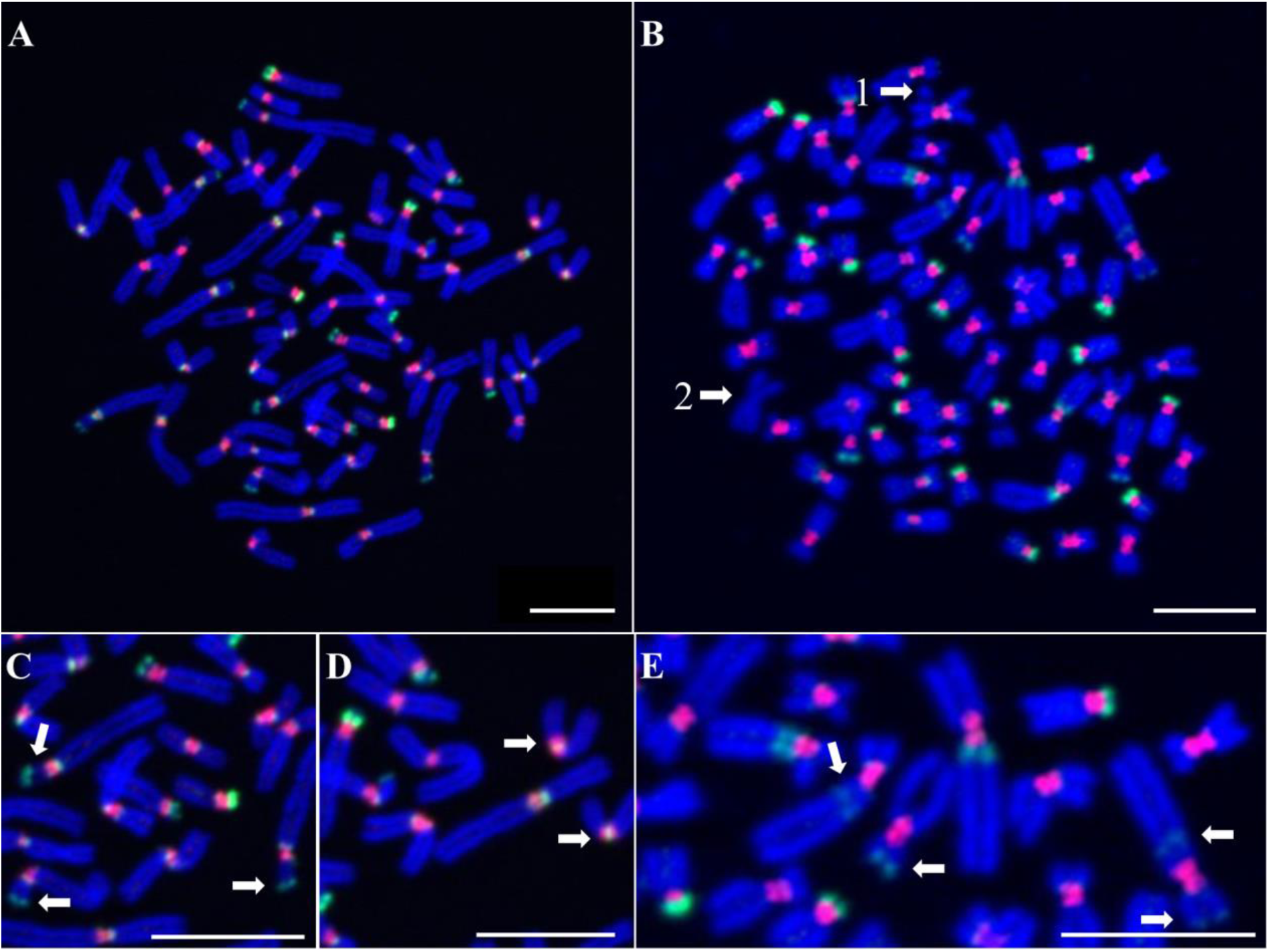
FISH analysis of the C1 and C2 alpha satellite families in species from the arboreal clade. Metaphase chromosomes are shown in blue (DAPI counterstain). Hybridization signal of probes C1b (red) and C2b (green) on: (A) Chromosomes of a female *C. roloway*, and (B) Chromosomes of a male *C. cephus*. Arrow 1 points to the unlabeled Y chromosome. Arrow 2 points to one autosome displaying a complete absence of signal for both C1 and C2 families. (C) Magnified view of image (A). Arrows indicate C2 signals observed in telomeric regions. (D) Magnified view of image (A). Arrows indicate two chromosomes where C1 signals (red) are localized in the pericentromeric region, surrounding the C2 signal (green) located in the core centromere. (E) Magnified view of image (B). Arrows indicate C2 signals observed in telomeric regions or in interstitial positions within the long arms. Scale bar = 10 µm.

C5 probe signals were observed in all species of the arboreal clade, but displayed distinct distribution patterns correlating with karyotype structure (Fig. 4). In the species possessing the least fissioned karyotypes (*C. roloway* and *C. diana*), C5 signals were restricted to the core centromeres of 6 chromosome pairs and did not extend toward the pericentromeres (Fig. 4A, D). Conversely, in species with highly fissioned karyotypes (*C. mona*, *C. cephus*, *C. erythrotis*, *C. ascanius*, *C. nictitans* and *C. mitis*), C5 signals extended toward the pericentromeres and were exclusively pericentromeric on several chromosomes (Fig. 4B, E). All C5 signals were persistent after stringent washing conditions. C6 probe signals were detected exclusively at core centromeres in the species with highly fissioned karyotypes. In particular, substantial signals were observed on acrocentric chromosomes in all these species (Fig. 4C).

#### Swamp clade of Cercopithecini

The representative species of the swamp clade in the present study is *A. nigroviridis* (2n=48). In *A. nigroviridis*, the distribution of alpha satellite families reveals a distinctive pattern when compared to other Cercopithecini species (Figs. 5 and S6).

Signals were observed with C1 probes on several chromosomes. However, the two C1 probes employed (C1a and C1b, Table S4), which target different sites on C1 sequences, revealed non-overlapping signals (Fig. S6). This lack of co-localization indicates that the sequences detected on *A. nigroviridis* chromosomes do not belong to the canonical C1 family. Consequently, these results imply the absence of the C1 family from the *A. nigroviridis* genome.

Clear FISH signals for the C2 family were observed on the core centromeres of all chromosomes, with the single exception of the Y chromosome. Moreover, the C2 family was also detected on the extremity of the short arms of the acrocentric chromosomes (Fig. 5A, D). The C3 and C4 probes provided clear signals with a distribution highly similar to C2, localizing to the core centromeres of all chromosomes (except the Y chromosome) and to the extremity of the short arms of the acrocentric chromosomes (Fig. 5B, C, E). Notably, a unique chromosome was specifically labeled by C3 probes but showed no hybridization with C4 probes (Fig. 5E).

**Figure 4.**
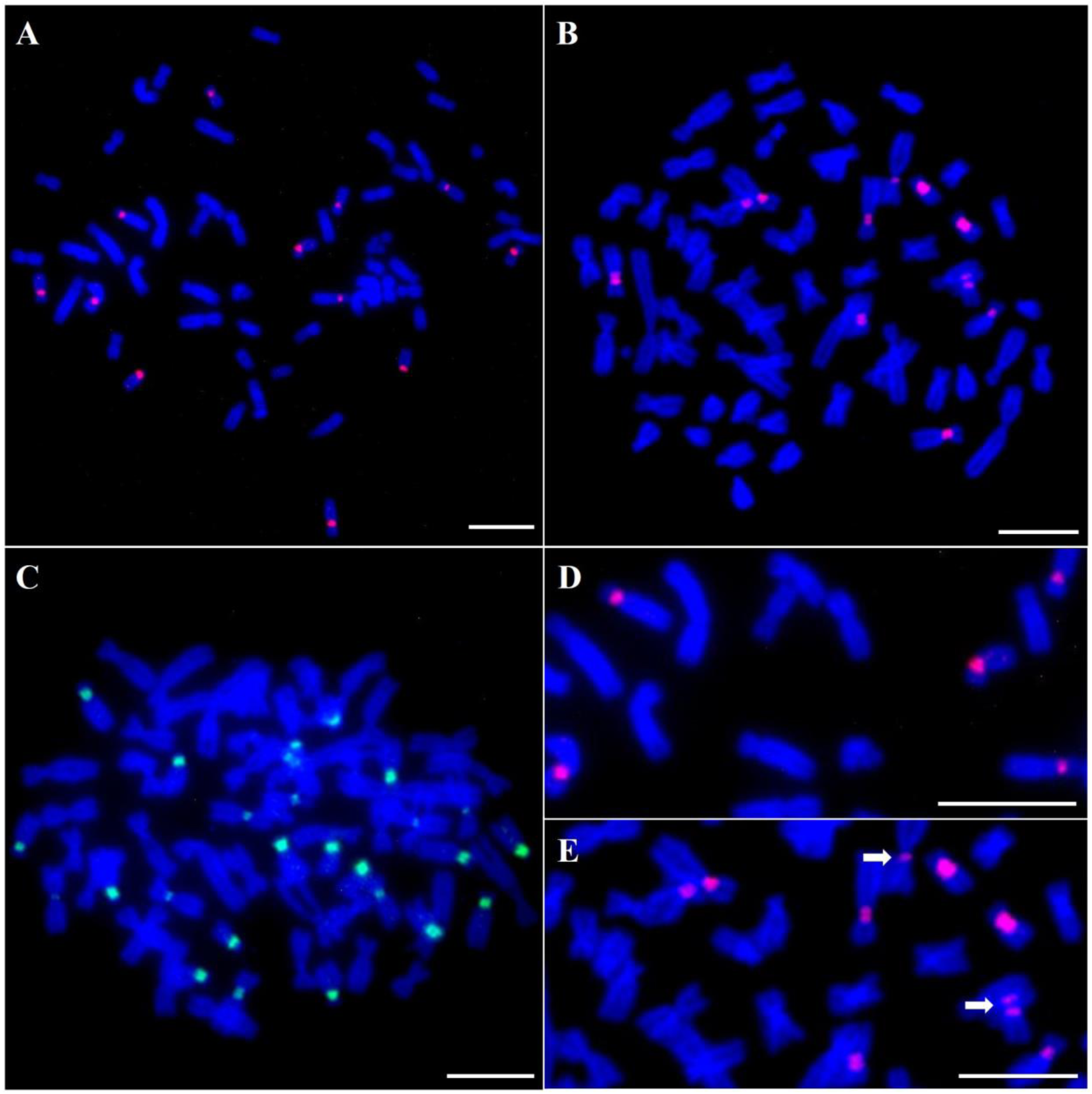
FISH analysis of the C5 and C6 alpha satellite families in species from the arboreal clade. Metaphase chromosomes are shown in blue (DAPI counterstain). Hybridization signal of probe C5a (red) on: (A) Chromosomes of *C. roloway*, and (B) Chromosomes of *C. cephus*. (C) Hybridization of probe C6a (green) on *C. cephus* chromosomes. Substantial signals are observed on acrocentric chromosomes. (D) Magnified view of image (A). C5 signals (red) are localized exclusively to the core centromeres. (E) Magnified view of image (B). Arrows point to chromosomes where C5 signals (red) are localized exclusively in the pericentromeric region. Scale bar = 10 µm.

**Figure 5.**
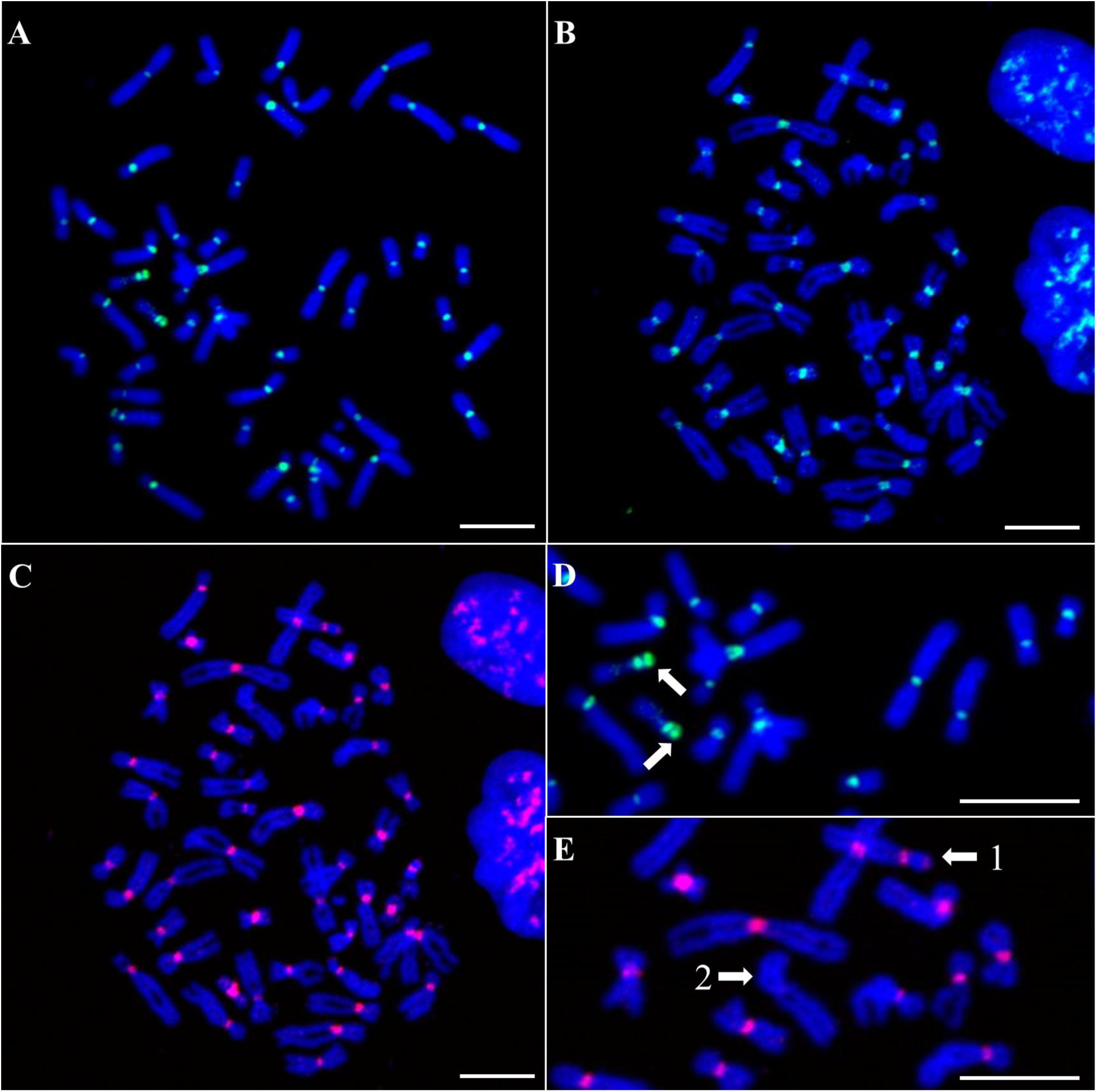
FISH analysis of the C2, C3 and C4 alpha satellite families in the swamp clade species *Allenopithecus nigroviridis*. Metaphase chromosomes are shown in blue (DAPI counterstain). (A) Hybridization signal of probe C2b (green). (B) Hybridization signal of probe C3a (green). (C) Hybridization signal of probe C4a (red). (D) Magnified view of image (A). Arrows point to labeled extremities of acrocentric chromosome short arms. (E) Magnified view of image (C). Arrow 1 points to the labeled extremity of an acrocentric chromosome short arm. Arrow 2 points to an unlabeled chromosome. Scale bar = 10 µm.

C5 probes produced slight signals on *A. nigroviridis* chromosomes, exhibiting a temperature-dependent signal reduction similar to that previously observed in *A. lhoesti*: the intensity of C5 signals was significantly reduced when the stringent post-hybridization wash was performed at 68 °C, compared to the standard temperature of 63 °C. This result suggests non-specific hybridization, leading to the conclusion that the C5 family is absent from the *A. nigroviridis* genome. C6 probes did not yield any detectable signals on the chromosomes of *A. nigroviridis*.

#### Papionini and Colobinae (Outgroups)

To contextualize the distribution patterns observed within Cercopithecini, FISH was performed on two outgroup species: *Macaca sylvanus* (Papionini) and *Colobus angolensis* (Colobinae).

The assessment of the C1 and C5 families in *M. sylvanus* provided patterns consistent with those observed in *A. nigroviridis*: probes C1a and C1b revealed non-overlapping signals, and C5 probes yielded signals that exhibited a temperature-dependent signal attenuation. These results indicate that the sequences detected are not homologous to the canonical C1 and C5 families, implying the absence of these alpha satellite families from the *M. sylvanus* genome.

Similarly, the analysis of the C2 family using two distinct probes (C2a and C2b, Table S4) resulted in non-overlapping signals in *M. sylvanus* (Fig. S7). This lack of co-localization indicates that the detected sequences are not homologous to the canonical C2 family, suggesting the absence of this alpha satellite family from the *M. sylvanus* genome.

Clear hybridization signals were observed for both the C3 and C4 probes, localizing to the centromeric regions in *M. sylvanus* (Fig. S8). However, the signals generated by the C3 and C4 probes did not exhibit a clear co-localization. This observation suggests that, unlike the pattern identified in Cercopithecini, the C3 and C4 families are not organized as obligate dimeric C3-C4 sequences in *M. sylvanus*.

Finally, hybridization signals for C6 probes were detected across the centromeres of all *M. sylvanus* chromosomes. Given that C6 was phylogenetically identified as a derived C1 subfamily (Cacheux et al., 2018) and had, until now, been detected only in a restricted subset of arboreal Cercopithecini characterized by highly derived karyotypes, its detection in the Papionini species *M. sylvanus* was unexpected. Considering the ancient divergence between Cercopithecini and Papionini and the absence of documented recent introgression, this observation likely reflects sequence convergence rather than shared ancestry; specifically, C6 probes likely detect a distinct alpha satellite family in *M. sylvanus* that coincidentally possesses the target sequence.

In the second outgroup species, *C. angolensis*, hybridization signals were detected exclusively with C5 probes. However, the intensity of these signals decreased substantially when the post-hybridization wash temperature was raised from the standard 63 °C to 68 °C. This temperature-dependent signal reduction is indicative of non-specific hybridization, allowing us to conclude that none of the C1 to C6 alpha satellite families are present in the *C. angolensis* genome.

### Detailed distribution of alpha satellite families on Cercopithecini chromosomes

The comparative analysis of the six representative Cercopithecini species focused on mapping the C2, C5, and C6 alpha satellite families. The C1 family was excluded from the detailed mapping because, where present, it showed minimal inter-species variation in centromeric localization. The C3/C4 families were excluded because their hybridization signals were either absent (below the FISH detection limit) or consistently too faint to allow for accurate centromeric localization across most species. The detailed distribution profiles for C2, C5, and C6 are presented in the schematic alignment of homologous chromosomes (Fig. 6), which includes the presumed ancestral Cercopithecidae karyotype (ANC). This schematic representation was generated by mapping the distribution data from the completed FISH karyotypes (Figs. S9, S10, and S11) onto the underlying structure for homology provided by the R-banding alignment (Fig. S12). FISH data for *A. solatus* and *C. pogonias* were retrieved from previous studies (Cacheux et al., 2016, 2018).

#### Inter-species homologous chromosomes

C2 probes yielded strong FISH signals on the short arms of all acrocentric chromosome pairs across all species investigated (Fig. 6 and Fig. S9). In several instances, substantial C2 signals were detected in the centromeric regions of homologous chromosomes. This is evident, for example, on the homologs of ANC4 (ANI2, EPA2, CRO1, and CCE1) (Fig. 6). Furthermore, C2 signals appeared in the telomeric regions of homologs to ANC9 (EPA10, CRO9, and CCE4).

C5 signals were also detected in the centromeric regions of homologous chromosomes, such as the homologs of ANC1 (CRO21, CCE18, and CPO15) (Fig. 6 and Fig. S10). In contrast, some chromosomes displayed C5 signals exclusively in single species, exemplified by CRO9 (homolog to ANC9). C5 signals were observed at the centromere of all acrocentric chromosomes in *E. patas*, but only on a single acrocentric pair in *C. roloway* (ANC1: CRO26). It should be noted that C5 signals were present on only one chromosome of the EPA2 pair (ANC4) in the studied specimen.

C6 signals were detected at the centromere of all acrocentric chromosomes in both *C. cephus* and *C. pogonias* (Fig. 6 and Fig. S11). A majority of these loci are homologous between the two species. In addition, C6 signals were observed on two homologous submetacentric chromosomes: CCE9 and CPO8 (ANC11). Fainter C6 signals were sporadically observed on non-homologous meta- or submetacentric chromosomes, such as CPO12 (ANC5), CPO11 (ANC15), and CCE19 (ANC18). Lastly, C6 signals were detected on a single chromosome of the CPO2 pair (ANC6) in the studied specimen.

#### Inter-species homologous centromeres

As detailed previously, C2 and C5 probe FISH signals showed variable localization, mapping either to the centromeres or pericentromeres depending on the species investigated. We found that strong C2 signals were present at the core centromere of *A. nigroviridis* chromosomes but localized to the pericentromeres in some homologous chromosomes of other species, such as the homologs of ANC4 (ANI2, EPA2, CRO1, and CCE1) (Fig. 6 and Fig. S9).

Similarly, C5 signals were observed at the core centromere of *E. patas* and *C. roloway* chromosomes, while localizing to the pericentromeres of some other homologs. See, for instance, the homologs of ANC6 (CRO10, CCE5, and CPO2) or ANC11 (EPA8, CCE9, and CPO8) (Fig. 6 and Fig. S10).

#### Putative evolutionary new centromeres

Chromosomes possessing a putative Evolutionary New Centromere (ENC) are indicated by black stars in Figure 6. Based on banding patterns, these chromosomes are homologous to presumed ancestral chromosome arms and are hypothesized to be the result of non-centromeric fissions of ancestral chromosomes (Dutrillaux et al., 1980, 1982; Moulin et al., 2008; see Fig. S12). Strong FISH signals provided by C2 probes were observed at the centromeric region of a large proportion of these putative ENCs. C2 signal detection on putative ENCs varied across species (FISH-positive loci/total putative ENCs): 2/2 for *A. nigroviridis*, 2/3 for *E. patas*, 3/6 for *A. solatus*, 6/7 for *C. roloway*, 8/11 for *C. cephus*, and 11/14 for *C. pogonias* (Fig. 6 and Fig. S9).

In contrast, C5 signals were detected only sporadically on putative ENCs, specifically 2/3 for *E. patas*, 2/7 for *C. roloway*, and 1/11 for *C. cephus* (Fig. 6 and Fig. S10).

Finally, C6 signals were observed on most of the putative ENCs in *C. cephus* (8/11) and *C. pogonias* (11/14), combined with pericentromeric C2 signals (Fig. 6 and Fig. S11). All putative ENCs detected with C2 and C6 signals were located on acrocentric chromosomes (CCE24-27, CCE29-32; CPO24-27, CPO29-35). CCE28 and CPO28 were the only acrocentric chromosomes that did not possess a putative ENC in *C. cephus* and *C. pogonias*, but still displayed the C2/C6 combination.

**Figure 6.**
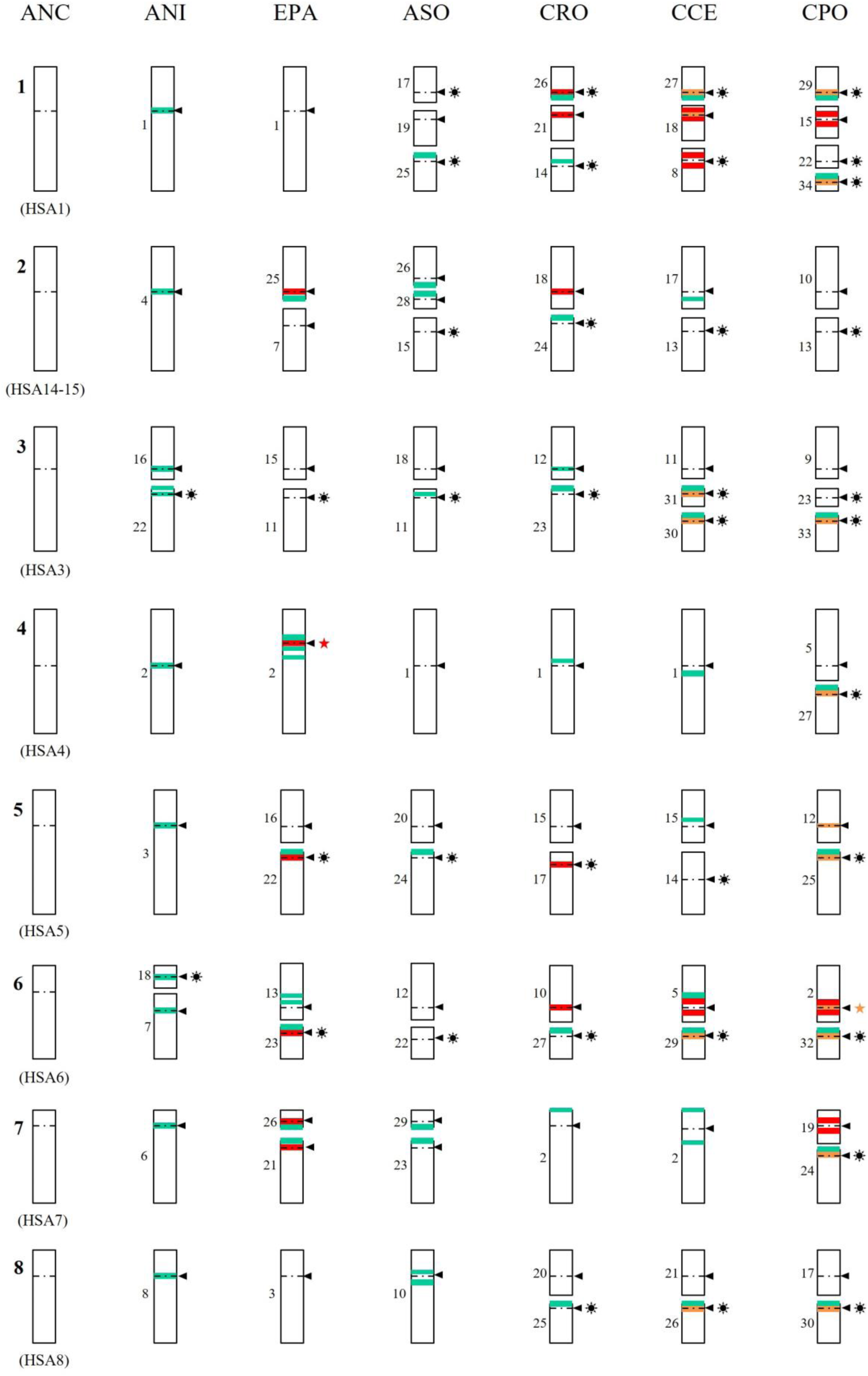
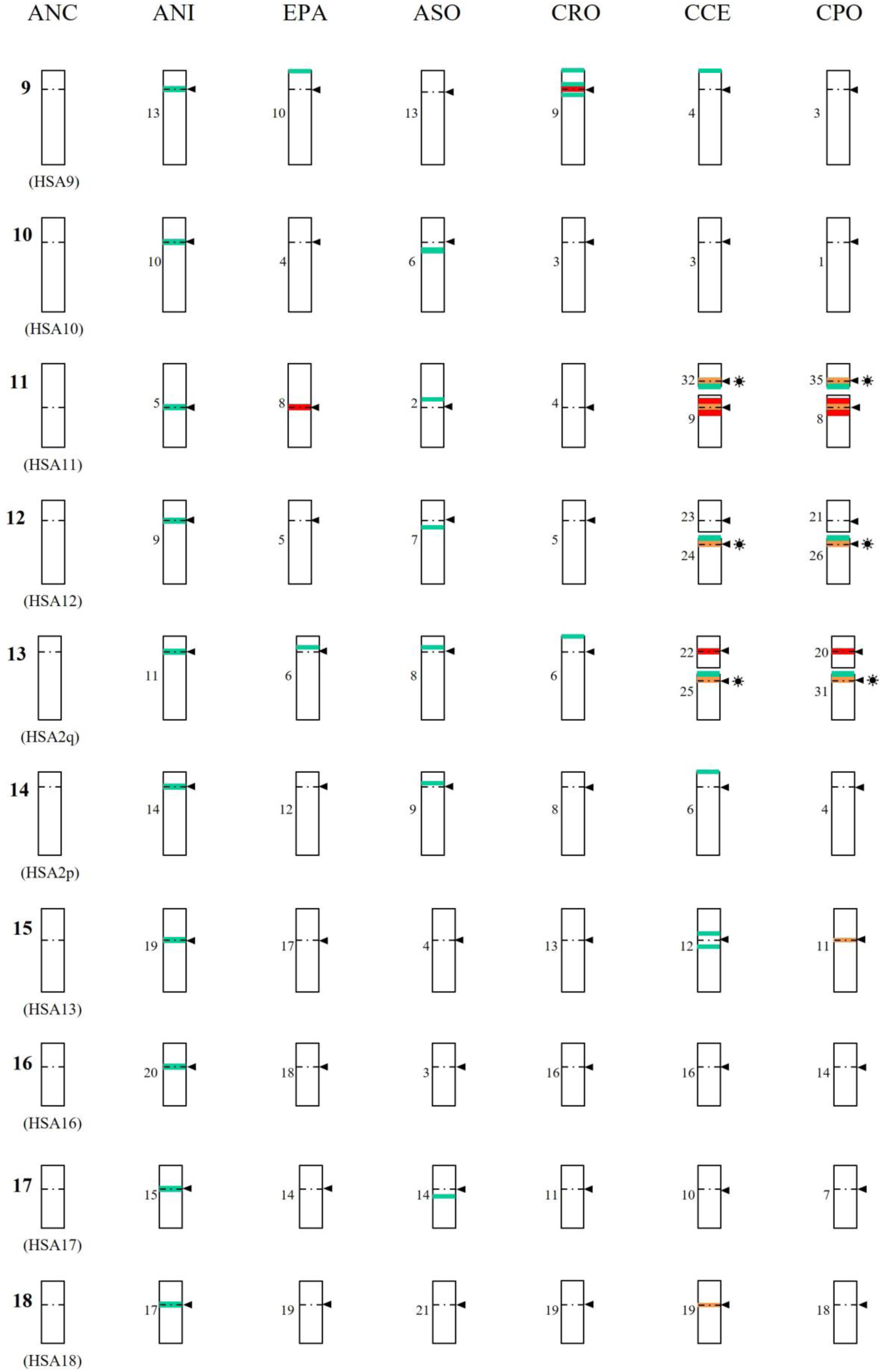
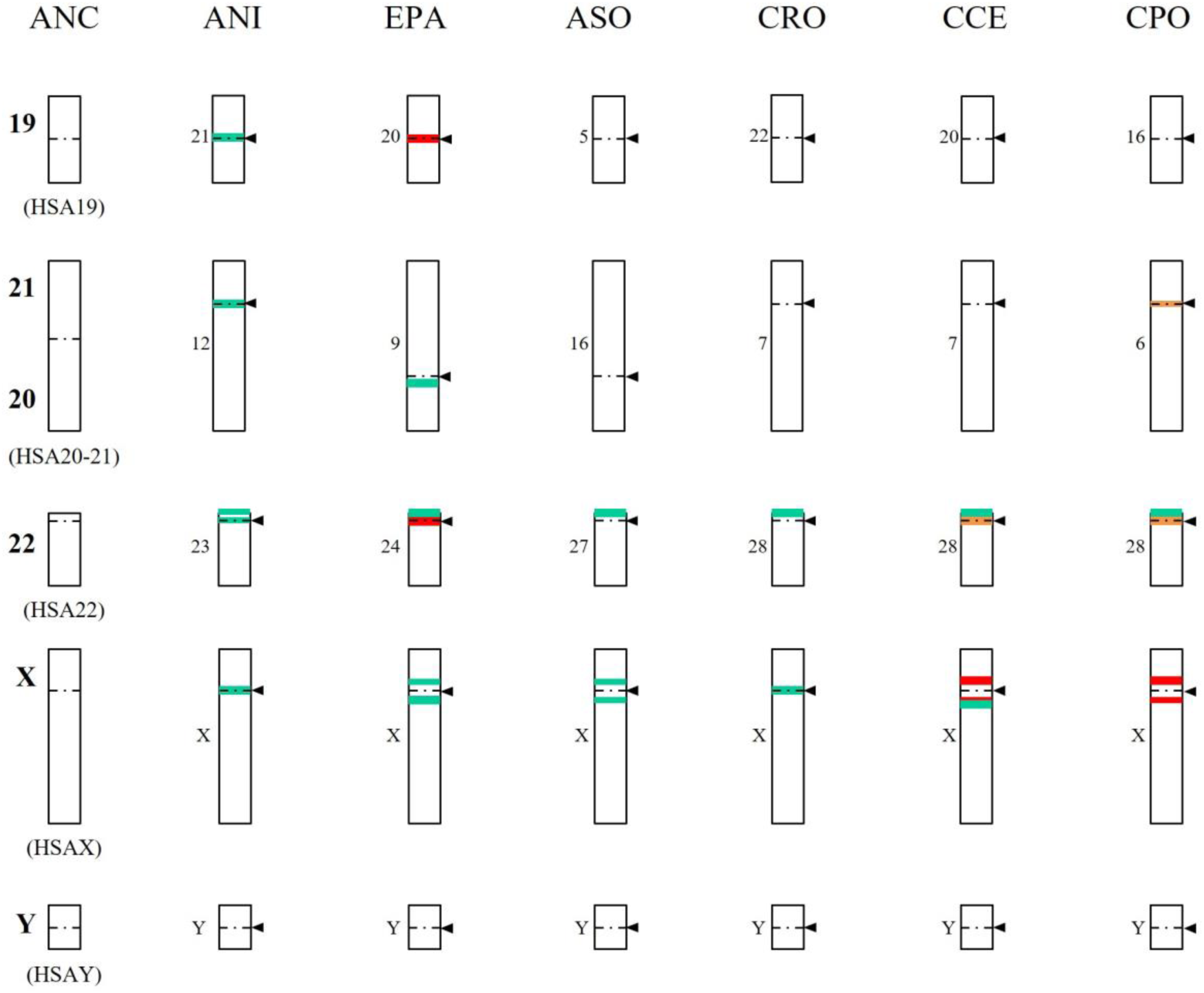
Schematic representation of C2, C5, and C6 alpha satellite distribution on homologous chromosomes in Cercopithecini. This schematic illustrates the alignment of homologous chromosomes across six species: *A. nigroviridis* (ANI), *E. patas* (EPA), *A. solatus* (ASO), *C. roloway* (CRO), *C. cephus* (CCE), and *C. pogonias* (CPO), with the presumed ancestral Cercopithecidae karyotype (ANC) serving as the reference structure. The chromosomal alignment is based on published R-banding patterns and chromosome painting data (Dutrillaux et al., 1982; Moulin et al., 2008). Chromosome numbers (in reference karyotypes) are displayed on the left side (Dutrillaux et al., 1978, 1980, 1988; Moulin et al., 2008). Homologies with human chromosomes (HSA) are mentioned. The distribution data for the alpha satellite families were mapped onto this structure using the completed FISH karyotypes (Figs. S9-S11). Alpha satellite distribution is shown in pastel green for C2, red for C5, and orange for C6. **C2 Visualization Criterion**: Only centro- or pericentromeric loci displaying the strongest FISH signals are illustrated (C2 probes provided signals of varying intensity on almost every chromosome). In contrast, all telomeric or interstitial loci exhibiting C2 signals are systematically included. **Symbols**: Arrows and dotted lines indicate centromere positions. Black stars denote putative evolutionary new centromeres. Colored stars indicate heterozygosity, where C5 (red) or C6 (orange) signals were observed on a single chromosome of the concerned pair.

## Discussion

In the present study, we combined classical and molecular cytogenetics in a comparative study to comprehensively map the distribution of six alpha satellite families (C1 to C6) across the chromosomes and centromere regions of 13 Cercopithecini species. We reconstructed a molecular phylogeny using nuclear markers to interpret our results within an evolutionary framework. This phylogeny resolves three distinct Cercopithecini clades: the established terrestrial and arboreal lineages, which comprise species with highly fissioned karyotypes, and a newly designated ‘swamp clade’, which notably includes *Allenopithecus nigroviridis*, a species characterized by a highly conserved karyotype. The presence and distribution pattern of each alpha satellite family on chromosomes and centromere regions varies significantly between lineages and even species. Notably, families previously described as heterogeneous in two terrestrial and arboreal species are present in all three clades. They exhibit a centromeric position in the swamp lineage but show a pericentromeric distribution in the terrestrial and arboreal lineages. Conversely, families previously described as homogeneous are strictly confined to the terrestrial and arboreal lineages, still showing variable centromeric/pericentromeric positioning that correlates with the degree of karyotype fissioning. Within the arboreal clade, including the species with the most highly-fissioned karyotypes, putative Evolutionary New Centromeres (ENCs) display a unique alpha satellite diversity, marked by the co-occurrence of the most homogeneous family and one of the most heterogeneous families identified.

### Tree-based alpha satellite evolutionary history in Cercopithecini

Reconstructing a phylogeny that encompasses our complete taxonomic sampling was essential to include all 13 Cercopithecini species studied by FISH, as some – notably *Cercopithecus erythrotis* – were absent from recent genome-wide datasets (Jensen et al., 2023, 2024). The molecular phylogeny we obtained is largely consistent with previous phylogenies based on nuclear (Tosi et al., 2004; Xing et al., 2007; Perelman et al., 2011; Jensen et al., 2023, 2024), chromosomal (Dutrillaux et al., 1980, 1982; Moulin et al., 2008), morphological (Cardini and Elton, 2008), and vocalization data (Gautier, 1988), particularly in resolving the established terrestrial and arboreal clades. Our phylogeny also delineates a new ‘swamp clade’ comprising *Allenopithecus* and *Miopithecus* species, which is positioned here as sister to the terrestrial and arboreal lineages. Notably, this swamp clade was not recovered in genome-wide analyses, which place *Allenopithecus* and *Miopithecus* as basal to the terrestrial and arboreal clades, respectively; such topological discordances are common in Cercopithecini and may reflect ancient hybridization or incomplete lineage sorting (Jensen et al., 2023, 2024).

Most nodes critical to reconstructing the alpha satellite evolutionary history in Cercopithecini are robustly supported in our phylogeny; however, one node exhibits lower statistical support: the node representing the common ancestor of the terrestrial and arboreal clades (PP = 0.70). Importantly, this specific node is further substantiated by the congruent distribution of alpha satellite families (as detailed below).

The monophyly of the clade grouping the Cercopithecini and the Papionini tribes, while beyond the scope of this study, has been extensively documented elsewhere (Page and Goodman, 2001; Rodrigues et al., 2009; Perelman et al., 2011).

### Evolutionary dynamics of alpha satellite DNA in Cercopithecini

Collectively, the findings from the present study and our previous work (Cacheux et al., 2016, 2018) allow us to propose a comprehensive evolutionary scenario for alpha satellite DNA dynamics within the Cercopithecini. This scenario, which illustrates the emergence and subsequent displacement of alpha satellite families, is summarized in Figure 1 and is anchored in time by phylogenetic estimations (Jensen et al., 2023).

#### C3 and C4 alpha satellite families

Sequencing experiments, coupled with subsequent *in silico* analyses (Cacheux et al., 2016; present study), revealed that the C3 and C4 families are associated into dimeric sequences within the *Allochrocebus solatus* and *C. pogonias* genomes. FISH experiments confirmed this association, showing colocalized C3 and C4 signals at the pericentromeres of *A. solatus* chromosomes (Cacheux et al., 2016). Notably, C3 and C4 signals were also detected on the *A. solatus* Y chromosome, a unique localization among the species studied so far. Conversely, in *C. pogonias*, the C3 family was largely undetectable, likely because its copy number has fallen below the FISH detection limit, while the C4 family was only faintly revealed at pericentromeres. This pattern of signal erosion is generalized across the terrestrial and arboreal clades, where only weak and infrequent signals for C3 and C4 were typically detected at pericentromeres. In sharp contrast, strong, clear signals were observed for both families at the centromeres of the swamp clade species, *A. nigroviridis*, and within the centromeric regions of the outgroup Papionini, *Macaca sylvanus*. Interestingly, C3 and C4 signals colocalized on *A. nigroviridis* chromosomes, yet showed non-overlapping patterns in *M. sylvanus*. Consequently, and consistent with the progressive proximal expansion model (Schueler et al., 2001, 2005), we propose that the C3 and C4 families emerged in the common ancestor of the Cercopithecinae (the taxonomic subfamily encompassing both the Cercopithecini and Papionini tribes) approximately 15 million years ago (Mya). Following this emergence, they amplified while remaining associated in Cercopithecini ancestors, dated to ∼13 Mya. Conserved at the centromere core in swamp clade species, C3 and C4 were subsequently displaced toward the pericentromeres in the common ancestor of the terrestrial and arboreal clades, estimated to be between 13 and 10 Mya. These older families now appear to be nearing extinction in the terrestrial and arboreal lineages, often exhibiting copy numbers too low to be reliably detected via FISH, and may totally disappear as newer families emerge in the genomes. Thus, the lifespan of alpha satellite families may not be equivalent across all evolutionary lineages, and the molecular mechanisms responsible for the erosion of satellite sequences at the pericentromeric borders have yet to be characterized.

#### C2 alpha satellite family

The C2 family was detected at the centromeres of the swamp clade species *A. nigroviridis* and was found at the pericentromeres of all terrestrial and arboreal clade species. This positional shift suggests that the C2 family probably emerged in a common Cercopithecini ancestor and was then displaced toward the pericentromeres in the common ancestors of the terrestrial and arboreal clade species. In addition, we observed the C2 family in the telomeric regions of certain submetacentric chromosomes across all three Cercopithecini lineages. Analogous instances of ectopic locations have been observed in other primate lineages, as alpha satellite sequences have been documented at the telomeric regions of two Hylobatidae species: the white-cheeked gibbon (*Nomascus leucogenys*) and the siamang (*Symphalangus syndactylus*) (Cellamare et al., 2009; Koga et al., 2012).

#### C1 alpha satellite family

The C1 family was not detected in *A. nigroviridis*. Conversely, strong C1 signals were observed at almost every centromeric regions of all terrestrial and arboreal clade species. The C1 family likely emerged in the common ancestor of the terrestrial and arboreal clade species, rapidly colonizing centromeres. This core family subsequently extended toward the pericentromeres while diversifying into the C5 and C6 subfamilies.

#### C5 alpha satellite family

The species and chromosomal distribution of the C5 family is notably more complex to interpret. This family was observed in all arboreal clade species and in a single terrestrial clade species, *Erythrocebus patas*. Two evolutionary scenarios may explain this observation.

Hypothesis 1: The C5 family may have emerged in the common ancestor of the terrestrial and arboreal clades, and subsequently undergone complete extinction in the terrestrial lineage leading to *Allochrocebus* species. However, this hypothesis contrasts with the evolutionary dynamics inferred for the C3 and C4 families, which are thought to have emerged ∼15 Mya and persisted ever since, leaving vestigial traces in most Cercopithecini genomes. In light of this long-term maintenance, the complete disappearance of C5 within a shorter period, without leaving any detectable pericentromeric traces, is plausible yet seems unlikely.

Hypothesis 2: Alternatively, the C5 family may have emerged solely in arboreal clade ancestors and been transferred to *E. patas* ancestors following genetic introgression. This hypothesis is supported by previous molecular studies that have suggested ancient hybridization events between arboreal clade lineages and *E. patas* ancestors (Guschanski et al., 2013; Jensen et al., 2023).

Regarding its localization, the C5 family adopts a centromeric position in *C. diana* and *C. roloway* but exhibits extension into pericentromeric regions in all other arboreal clade species. This pattern suggests that the C5 family initially emerged at the centromere in arboreal clade ancestors, estimated at ∼10 Mya, and was subsequently displaced toward the pericentromeres in the *C. mona* group ancestors and in the common ancestors of the *C. mitis* and *C. cephus* groups.

#### C6 alpha satellite family

Finally, the C6 family was exclusively detected at the centromeric core of species belonging to the *C. mona/mitis/cephus* groups. The C6 family thus represents the most recently emerged family identified in our studies, having arisen in the *C. mona* group ancestors and in the common ancestors of the *C. mitis* and *C. cephus* groups less than 10 Mya, and having been conserved at the centromere core ever since.

Importantly, the proposed scenario of alpha satellite family turnover not only aligns with the progressive proximal expansion model for alpha satellite DNA evolution (Schueler et al., 2001, 2005) but is also strongly supported by the observed sequence identities of each family. Under the common assumption that sequence identity correlates negatively with family age (i.e., younger families exhibit higher identity), our established successive emergence order for alpha satellite families in Cercopithecini is fully consistent with our sequence data. The average sequence identities observed are: C3: 87%; C4: 85%; C2: 85%; C1: 95%; C5: 95%; and C6: 98% within the *C. pogonias* genome (Cacheux et al., 2018; present study), and C3: 86%; C4: 83%; C2: 85%; and C1: 95% within the *A. solatus* genome (Cacheux et al., 2016). Consequently, the earlier classification of these families as either heterogeneous or homogeneous, based on sequence analysis in *A. solatus* and *C. pogonias*, appears to precisely reflects their respective early or late emergence during Cercopithecini evolution.

In this regard, sequencing and analyzing the alpha satellite DNA of *A. nigroviridis* would be highly relevant. This approach would help establish whether the sequence identities of the older families (C2, C3, and C4) are more homogeneous in *A. nigroviridis*, where these families appear conserved in the centromere core, than in species where they have been displaced toward the pericentromeres. A similar comparative analysis is warranted for the C5 family: specifically, investigating whether C5 exhibits greater sequence homogeneity in *C. diana* and *C. roloway* compared to *C. pogonias*, given its more centromeric distribution in the former two species.

Finally, when evaluating the validity of the proposed evolutionary scenario for alpha satellite families, it is essential to consider the inherent methodological limitations regarding both the exhaustiveness of family characterization and the subsequent cytogenetic analysis. As previously mentioned, our initial enzymatic isolation and sequencing of alpha satellite monomers may be non-exhaustive, potentially missing monomers that lack the restriction site of the enzyme utilized (Cacheux et al., 2016, 2018).

Furthermore, our two-species sampling strategy (*A. solatus* and *C. pogonias*) means lineage-specific alpha satellite families may exist in Cercopithecini species that were not included in our *in silico* characterization. This characterization bias is directly inherited by the present cytogenomic analysis. Indeed, discriminatory probes can only be designed from the sequences of known families: any families not retrieved by the enzymatic isolation step cannot be taken into account. Consequently, it is possible that some probes designed to specifically target an identified family may inadvertently target other families absent from our *in silico* datasets. A mitigation strategy was employed to address this limitation: whenever feasible, two independent probes were designed to target the same family at different sites of the constitutive monomers (Cacheux et al., 2016, 2018; this study). The co-localization of their respective signals strongly supports the specific detection of the intended family. Conversely, a non-overlapping signal suggests the detection of at least two distinct families, which are both different from the intended target family, as exemplified by the probes C1a and C1b in *A. nigroviridis*. If such probes reveal only a partial overlap of their respective signals, it can be hypothesized that the target family is distributed exclusively across the overlapping loci. Importantly, sequencing the alpha satellitome of additional Cercopithecini species beyond *A. solatus* and *C. pogonias* will be crucial for achieving a more complete characterization of alpha satellite families and obtaining an exhaustive view of their evolutionary dynamics.

### Centromere DNA evolution and chromosome rearrangement dynamics

The progressive proximal expansion of alpha satellite DNA at centromeres has traditionally been examined separately from the broader context of structural genome evolution. As a result, the impact of large-scale chromosome rearrangements on alpha satellite turnover remains unexplored in many primate lineages. However, the evolutionary history of the Cercopithecini reveals a compelling pattern: a remarkable correlation appears to exist between their active rearrangement dynamics – driven by frequent fissions – and the rapid turnover of their alpha satellite families. This observation suggests that the prevailing view, which separates centromere sequence evolution from genome structure evolution, may be incomplete and require re-evaluation.

We observe a clear contrast in the degree of rearrangement across the Cercopithecini clades since their divergence (Dutrillaux et al., 1980; Moulin et al., 2008). The swamp clade, exemplified by species like *A. nigroviridis* (2n = 48), exhibits a relatively stable karyotype which has conserved a structure close to the ancestral state, reflecting few evolutionary fissions since the divergence of the Cercopithecini. By contrast, the terrestrial clade shows a moderate increase in chromosome number (2n = 54 to 60) due to several fissions, and the arboreal clade harbors the most highly rearranged karyotypes, particularly in the *C. mona/mitis/cephus* groups (2n = 66 to 72), contrasting with the *C. diana* group (2n = 58). This dramatic divergence in rearrangement dynamics is tightly mirrored by the emergence and displacement of alpha satellite families. Specifically, the successive formations of novel centromeric arrays – such as C1 (in terrestrial/arboreal clade ancestors), C5 (in arboreal clade ancestors), and C6 (in ancestors of the *C. mona/mitis/cephus* groups) – are chronologically correlated with periods of major chromosome rearrangement dynamics. Similarly, the pericentromeric displacement of arrays C2, C3, C4 (in terrestrial/arboreal clade ancestors) and C5 (in ancestors of the *C. mona/mitis/cephus* groups) further signs these periods of significant karyotype evolution. This suggests that the evolutionary history of centromere sequence turnover is deeply linked to the structural evolution of the Cercopithecini genomes.

This relationship raises a pivotal question regarding the potential mutual causality between evolutionary turnover of alpha satellite families and chromosomal rearrangements in Cercopithecini. While chromosomal instability is often viewed as a consequence of cellular stress (e.g., Macheret and Halazonetis, 2015), it is equally important to consider that major evolutionary rearrangements (such as fissions) may themselves become chronic generators of cellular stress in the germline. We propose that chromosomal fissions generate a constellation of structural defects – including, but not limited to, the challenge of de novo telomere capping – thereby constituting a source of persistent cellular stress in the germline (e.g., Fagagna et al., 2003; Cesare and Reddel, 2010). This stress, in turn, may lead to the epigenetic deregulation and subsequent activation of transposable elements (TEs) (e.g., Fedoroff, 2012). Given the known propensity of many TEs to integrate preferentially into heterochromatic and pericentromeric regions, their mobilization could effectively destabilize centromere function (Ugarkovi¢, 2009; Plohl et al., 2014), creating a strong selective pressure for the emergence of new, stable, and highly homogeneous alpha satellite arrays. In this scenario, the evolutionary chromosomal rearrangements in Cercopithecini establish a permissive state that accelerates the centromere turnover, suggesting that karyotype structure and centromere sequence evolution are deeply intertwined.

### Diversity of alpha satellite DNA on evolutionary new centromeres

The increase in chromosome number observed during Cercopithecini evolution is thought to be driven largely by non-centromeric fissions (Dutrillaux et al., 1980; Stanyon et al., 2005; Moulin et al., 2008). These structural rearrangements necessitate the *de novo* formation of ENCs on the resulting acentric fragments to ensure their stable segregation and persistence across generations. The assembly of centromeric DNA sequences, including those associated with ENCs, remains exceptionally challenging, even with specialized sequencing and computational methodologies (Rocchi et al., 2012; Miga et al., 2020; Logsdon et al., 2024, 2025). Following the prediction by Stanyon et al. (2012), our combined approach of high-throughput sequencing and molecular cytogenetics allowed us to provide an integrated molecular characterization of the putative ENCs in several Cercopithecini species. Specifically, in the highly fissioned genomes of the arboreal clade species *C. pogonias* and *C. cephus*, most putative ENCs were found to be spanned by the newly emerged C6 alpha satellite family. Notably, these putative ENCs are systematically located on acrocentric chromosomes. The single exception is one homologous pair of acrocentric chromosomes (CPO28 and CCE28) that do not appear to be derived from a fission event: however, C6 is still detected at their centromeres, providing further evidence for peculiar inter-centromeric DNA exchanges between acrocentric chromosomes (Choo et al., 1990; Warburton et al., 2008; Cacheux et al., 2016, 2018). This hypothesis of exchange is further strengthened by the observation of the older C2 alpha satellite family on the short arm of all acrocentrics. We propose that C2 began colonizing these acrocentric chromosomes following exchange events originating from the older pericentromeres of CPO28 and CCE28.

Finally, it is important to acknowledge that the existence of ENCs in Cercopithecini, while strongly inferred from comparative banding patterns, remains to be definitively confirmed. This inference is particularly challenging given that apparent centromere shifts between species might be due not only to true centromere repositioning (i.e., ENCs) but also to pericentric inversions. Distinguishing centromere repositioning from inversions, especially those where the breakpoints are cryptic or undetectable using banding techniques, necessitates the use of molecular markers which reveal the order of flanking sequences relative to the centromere (Rocchi et al., 2012). Future investigations should prioritize high-resolution methods, such as BAC-FISH mapping (Ventura et al., 2001; Locke et al., 2011; Chiatante et al., 2016) or, ideally, comparative whole genome sequencing and assembly, to fully validate the de novo centromere status of the C6-spanned loci.

### Alpha satellite DNA as a tool to resolve phylogenetic relationships

While molecular phylogenetic studies often rely on coding or non-coding single-copy genes, it has been previously proposed that alpha satellite DNA could serve as a complementary molecular marker to resolve phylogenetic relationships (Shepelev et al., 2009). Our results support this concept by establishing distinct alpha satellite profiles that not only reflect major Cercopithecini clades, but also provide resolution for groupings that were either weakly supported or even not recovered in our gene phylogeny: 1) The C1 alpha satellite family was not detected in *A. nigroviridis* but was observed at almost every centromeric region of all terrestrial and arboreal clade species. Conversely, the C2, C3, and C4 families are retained at the centromeres of *A. nigroviridis*, but only persist as residual pericentromeric traces across terrestrial and arboreal species. This stark pattern strongly supports the existence of a clade comprising terrestrial and arboreal lineages, to the exclusion of the swamp lineage; 2) C6 was exclusively detected at the centromeric core of species belonging to the *C. mona/mitis/cephus* groups. This restricted distribution, combined with the chromosomal distribution of C5 in the arboreal clade species, supports the existence of a clade encompassing these three groups.

The phylogenetic implications of these alpha satellite profiles, however, may vary depending on the chosen evolutionary framework. Crucially, the external position of *A. nigroviridis* relative to the terrestrial/arboreal clade is supported by its unique alpha satellite signature; yet, this placement is incongruent with genome-wide analyses which position *Allenopithecus* as sister to the terrestrial clade (Jensen et al., 2023, 2024). This discrepancy leads to two alternative interpretations.

Interpretation 1: Centromeric introgression within the species tree. If the genome-wide phylogeny (Jensen et al., 2023, 2024) accurately represents the species’ history, *Allenopithecus* is sister to the terrestrial clade. In this case, the shared presence of C1 in both terrestrial and arboreal lineages – and its absence in *A. nigroviridis* – would imply an ancestral centromeric introgression between the exclusive ancestors of these two clades. Here, alpha satellite DNA would track a specific reticulate event that occurred after the divergence of *Allenopithecus*, rather than a vertical branching pattern.

Interpretation 2: Centromeric preservation of an ancestral signal. Alternatively, the alpha satellite distribution may reflect a deeper phylogenetic signal (also preserved in our multi-locus topology). In this view, the centromeres of *A. nigroviridis* would have retained an ancestral signature of the swamp lineage’s early divergence. Because centromeric regions largely escape meiotic recombination, they could have effectively bypassed the subsequent introgression and gene flow events that reshaped the rest of the nuclear genome, thereby preserving an original phylogenetic signal that has been elsewhere obscured (Mahtani and Willard, 1998; Talbert and Henikoff, 2010).

The support for a clade comprising the *C. mona/mitis/cephus* groups further illustrates the potential of alpha satellite DNA for resolving complex evolutionary histories, thereby clarifying the external position of the *C. diana* group within the arboreal clade. While the placement of the latter remains a subject of debate – often showing weak statistical support even in genome-wide datasets (Jensen et al., 2023, 2024) – this basal divergence is here corroborated by specific alpha satellite signatures, in addition to being supported by chromosomal data (Moulin et al., 2008). Alternatively, and as previously discussed, the shared alpha satellite profile between the *C. mona/C. cephus/C. mitis* groups could also be interpreted as the result of an ancestral centromeric introgression between the *C. mona* group ancestors and the common ancestors of the *C. mitis* and *C. cephus* groups, providing a signature of reticulate evolution that goes beyond a strictly vertical phylogenetic signal.

## Conclusion

This work provides an integrated cytogenomic framework for elucidating the dynamic and interwoven histories of alpha satellite DNA and karyotype evolution across the Cercopithecini. Our results lend support to the progressive proximal expansion model while illuminating a coherent evolutionary scenario, encompassing the successive emergence of alpha satellite families at core centromeres, their eventual displacement toward pericentromeres, and their ultimate extinction. Crucially, our findings reveal an intimate correlation between centromere and chromosome rearrangement dynamics, which we interpret as a co-evolution between alpha satellite sequences and the overall architecture of the genome. Ultimately, the evolutionary history encoded within these repetitive elements offers a unifying concept: the genome as an ecosystem where the birth, diversification, and extinction of alpha satellite families mirror the fundamental evolutionary dynamics of biological species.

## Acknowledgments

The authors would like to thank Jérôme Fuchs for his valuable assistance and expertise with the phylogenetic analyses, as well as Julien Pichon for his careful reading of the manuscript and constructive comments.

## Data, scripts, code, and supplementary information availability

Data, input files, and supplementary information are available online in the Zenodo repository: https://doi.org/10.5281/zenodo.19650370. These include the concatenated sequence alignment, PartitionFinder configuration and results files, MrBayes command blocks, and the following supplementary information:

Figure S1: FISH analysis of the C3 and C4 alpha satellite families in species from the terrestrial clade.

Figure S2: FISH analysis of the C5 alpha satellite family in species from the terrestrial clade.

Figure S3: Classification of alpha satellite monomers, retrieved from Cercopithecus pogonias dimeric sequences, within the C3 and C4 families.

Figure S4: Inter-species comparison of alpha satellite monomers retrieved from Cercopithecus pogonias and Allochrocebus solatus dimeric sequences.

Figure S5: FISH analysis of the C4 alpha satellite family on Cercopithecus pogonias chromosomes.

Figure S6: FISH analysis of the C1 alpha satellite family on Allenopithecus nigroviridis chromosomes.

Figure S7: FISH analysis of the C2 alpha satellite family on Macaca sylvanus chromosomes.

Figure S8: FISH analysis of the C3 and C4 alpha satellite families on Macaca sylvanus chromosomes.

Figure S9: Distribution pattern of the C2 alpha satellite family on Cercopithecini chromosomes.

Figure S10: Distribution pattern of the C5 alpha satellite family on Cercopithecini chromosomes.

Figure S11: Distribution pattern of the C6 alpha satellite family on Cercopithecus cephus chromosomes.

Figure S12: Alignment of homologous chromosomes based on R-banding patterns across Cercopithecini species.

Table S1: PCR primers used for phylogenetic studies.

Table S2: GenBank sequences included in phylogenetic studies.

Table S3: Specimens from the TCCV, RBCell collection (MNHN) included in cytogenomic studies.

Table S4: Oligo-FISH probe sequences and associated labels.

Table S5: Summary of Oligo-FISH probe sets used per species.

Table S6: Observed counts of alpha satellite family associations in *Cercopithecus pogonias* dimeric sequences.

Table S7: Average sequence identities in alpha satellite families identified in Cercopithecini (C1-C6).

